# Evolution of sexually-transferred steroids in *Anopheles* mosquitoes

**DOI:** 10.1101/248112

**Authors:** Emilie Pondeville, Nicolas Puchot, Michael Lang, Floriane Cherrier, Francis Schaffner, Chantal Dauphin-Villemant, Emmanuel Bischoff, Catherine Bourgouin

## Abstract

Human malaria, which remains a major public health problem, is transmitted by a subset of *Anopheles* mosquitoes belonging to only three out of eight subgenera: *Anopheles, Cellia* and *Nyssorhynchus*. Unlike almost every other insect species, it was shown that males of some *Anopheles* species produce and transfer steroid hormones to females during copulation and that this transfer mediates reproductive changes. Steroids are consequently seen as a potential target for malaria vector control. Here, we analysed the evolution of sexually-transferred steroids and their effects on female reproductive traits across *Anopheles* by using a set of 16 mosquito species (5 *Anopheles*, 8 *Cellia*, and 3 *Nyssorhynchus*), including malaria vector and non-vector species. We show that male steroid production and transfer are specific to the *Cellia* subgenus and that there is no correlation between mating-induced effects in females and sexually-transferred steroids. In the light of our results, male steroid production, transfer and post-mating effects in females do not correlate with their ability to transmit human malaria, which overturns the suggestion from previous studies and suggests that manipulation of steroid-response pathways in the field should be considered with caution in order to benefit malaria vector control strategies.

## Introduction

*Anopheles* mosquitoes are mostly known for their ability to transmit to mammals malaria caused by *Plasmodium* parasites. Among 472 species named in this genus, 41 have been classified as dominant vector species (DVS) of human malaria [1–3]. The *Anopheles* genus is further subdivided into 8 subgenera of which 3 (*Anopheles, Cellia*, and *Nyssorhynchus*) contains all known DVS of human malaria [3]. Despite a significant decrease of malaria incidence in the last 15 years due to vector control and improvement of chemoprevention, diagnostic testing and treatment, human malaria remains a world-wide burden with more than 200 million cases reported and an estimated 445 000 deaths in 2016 [4]. With the increase of insecticide resistance in malaria mosquitoes [4, 5], new vector control strategies to limit and possibly eliminate malaria transmission are being developed such as replacement of wild malaria-susceptible mosquito populations by resistant ones and/or decrease of vector populations by manipulating mosquito reproduction [6–12]. Unlike vertebrates, it was considered that insect adult males do not produce significant amounts of steroid hormones until it was shown that males of *Anopheles gambiae*, the main vector of human malaria in Africa, produce and transfer high quantities of 20-hydroxyecdysone (20E) to females during copulation [13]. While ovarian steroid synthesis occurs in mosquito females upon blood meal triggering egg development [13–18], sexual transfer of steroids by males likely represents a nuptial gift that affects female reproduction in the malaria vector. Consistent with this, sexually-transferred steroids were shown to induce refractoriness to further copulation and to stimulate oviposition in *An. gambiae* females [19]. A study further reported that mating-induced phenotypes in *Anopheles* females would only occur in species whose males produce and transfer steroids to females during mating and that would be specific to dominant human malaria vectors. Since steroids contribute to promote oogenesis and possibly favouring *Plasmodium* survival, the authors concluded that sexual transfer of steroids is likely to have shape anopheline ability to transmit malaria to humans [20]. Steroids became as such a promising target to manipulate mosquito female reproduction with the aim to reduce malaria vector populations specifically. Consequently, field use of 20E receptor agonists has been recently proposed to reduce malaria transmission [21]. In the present work, we investigated the evolutionary history of male steroid production using a large set of mosquito species belonging to the three *Anopheles* subgenera *(Anopheles, Cellia, Nyssorhynchus)* that contain all known DVS of human malaria. We also investigated the post-mating effects on female reproductive traits whether the cognate males produce or not steroids. The main conclusions of our work are that i) production of steroids in male mosquitoes is restricted to the *Cellia* subgenus, and ii) the effect of mating on female reproductive potential vary across species of the 3 *Anopheles* subgenera and this is not correlated to male steroid production.

## Results

### Male steroid production is specific to the *Cellia* subgenus

We first measured steroid titers in sexually mature virgin males from 19 different mosquito species. These mosquito species were selected within two *Culicidae* subfamilies, *Anophelinae* and *Culicinae*. Within the *Anophelinae* subfamily, 16 species distributed all over the world were chosen to cover 3 different subgenera (*i.e. Cellia, Nyssorhynchus* and *Anopheles*) of the *Anopheles* genus (Figure 1A). In the *Culicinae* subfamily, *Aedes aegypti, Aedes albopictus* and *Culex pipiens* that are vectors of arboviruses were also analysed. As shown on Figure 1B (right panel), male 20E production occurs only within the *Cellia* subgenus (*Anophelinae*) and is absent in the *Culicinae*. Interestingly, *Anopheles quadriannulatus* whose males produce similar levels of steroids compared to *Anopheles stephensi* is not a malaria vector unlike *An. stephensi* and other *Cellia* species investigated in the present study. Indeed, its refractoriness or low susceptibility to the human malaria parasite has been experimentally determined [22, 23], confirming epidemiological data [24]. Conversely, production of 20E was undetected in male mosquitoes from the two other *Anopheles* subgenera, *Anopheles* and *Nyssorhynchus* of which all tested members are registered as DVS and/or experimentally shown to be highly susceptible to *Plasmodium falciparum* [25, 26].

**Figure 1.**
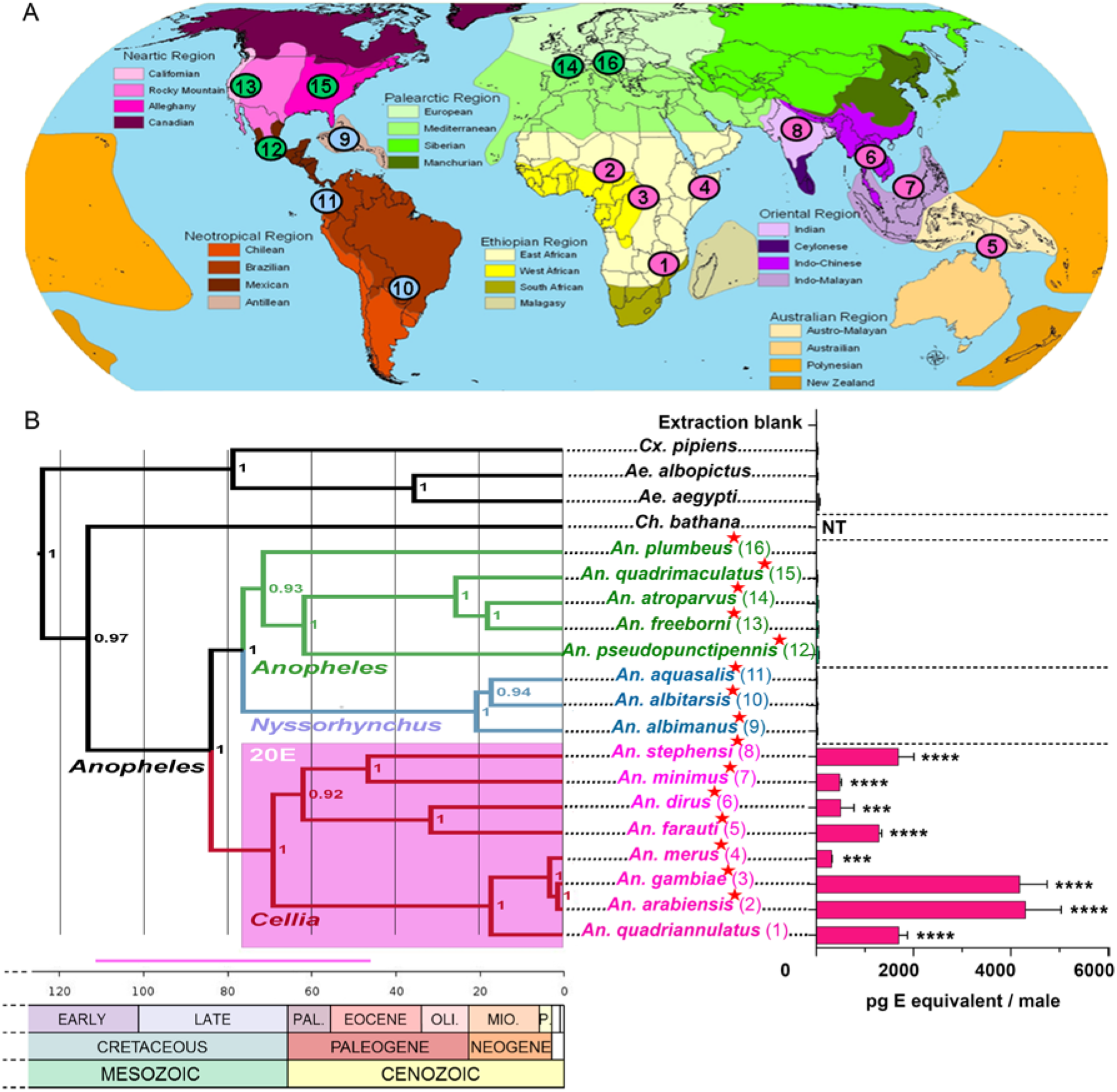
Distribution, phylogenetic relationships of mosquitoes and steroid production in mosquito adult males. (A) Present geographical distribution of the 16 mosquito species (*Anopheles* genus) analysed in this study represented on a zoogeographical map (modified from The geographical distribution of animals, Eckert IV projections department of geosciences, Texas tech university). Species matching numbers are shown on panel B. Pink: *Cellia* subgenus, blue: *Nyssorhynchus* subgenus, green: *Anopheles* subgenus. (B) Bayesian phylogeny of 20 *Culicidae* species, 19 species tested for ecdysteroid male production plus *Chagasia bathana* (subfamily *Anophelinae*, genus *Chagasia*) used as outgroup for phylogenetic analyses. Dominant malaria human vectors are indicated by a red star. Time is represented in millions of years (Ma). Approximated node ages are detailed in Supplementary Table 1. Bayesian node support values are presented on the right side of each node. Ecdysteroid titers in whole 5-day-old virgin males are indicated on the right side of the tree. Results are expressed as mean +/− SEM in pg E equivalents per male. Results were subjected to statistical analysis using Kruskall-Wallis test for nonparametric data followed by Dunn’s post-test (control group: Extraction blank). The indicated p values are those obtained with Dunn’s test (***, p value < 0.001; ****, p value < 0.0001). NT: not tested. Predicted lineages with significant male 20E production are shaded pink on the tree. The pink horizontal bar represents the minimum/maximum 95% confidence interval (CI) estimated time in Ma for origin of male 20E production. The geological time scale is adapted from the Geological Society of America (http://www.geosociety.org/science/timescale/). The white coloured cases represent the quaternary period. PAL., Paleocene; OLI., Oligocene; MIO., Miocene; P., Pliocene.

To uncover the evolutionary history of male steroid production in mosquitoes, we further consolidated the phylogenetic relationships of the species we analysed. To this effect, we used DNA sequences of partial regions of the coding sequence of mitochondrial genes (*COI, COII, ND5 and CYTB*) and nuclear genes (*g6pd* and *white* as well as ribosomal subunits *18S* and *28S*) from the 19 mosquito species plus *Chagasia bathana (Anophelinae* subfamily, *Chagasia* genera) as outgroup for *Anopheles* mosquitoes [27]. Phylogenetic analysis of the data set was performed by Bayesian inference (Figure 1B left panel) and by maximum likelihood (Supplementary Fig. 1). Phylogenetic relationships inferred from these analyses resulted in different topologies mainly at the subgenus level and with varying node support values. In both analyses, the members of each subgenus formed monophyletic groups and the branching orders within the subgenus *Cellia* was identical. Our results are in agreement with ones obtained by Sallum *et al*. [28] with the *Cellia* clade being the outgroup of the subgenera *Anopheles and Nyssorhynchus* in the Bayesian approach while in the maximum likelihood approach, *Nyssorhynchus* is the outgroup of *Anopheles* and *Cellia*, also in agreement with recently published phylogenies [1, 29, 30]. The *Anopheles* subgenus has also been found by others to be a basal lineage to *Cellia* and *Nyssorhynchus* [31]. Thus, relationships between the different subgenera *Cellia*, *Nyssorhynchus* and *Anopheles*, are still not entirely resolved. The difficulty in determining the relationships among these subgenera might be due to limits and bias in gene and taxon sampling, but also to different methods used [3, 27]. This could also be due to the fact that the radiation of *Anopheles* and *Cellia* species happened roughly at the same time [29, 31].

From the Bayesian phylogenetic analysis and using fossil data, we obtained species divergence estimates, which largely conform to recent phylogenies [29, 31, 32] (Supplementary Table 1). The age of the last common ancestor of *Anopheles* genus is about 84.1 Ma (112.7-55.8, 95% confidence interval) and the most ancestral node within the *Cellia* subgenus is dated to 69.2 Ma (93.0-45.6, 95% confidence interval) (Figure 1B, Supplementary Table 1). As males from all *Cellia* species tested so far have the ability to produce 20E, according to the parsimony law steroid production by mosquito males probably originated once in the early *Cellia* lineage, at about 84.1-69.2 Ma. (112.7-45.6, 95% confidence interval), *i.e*. during the late Cretaceous. Thus, steroid production by mosquito males is most probably a shared derived character from the last common ancestor of *Cellia* mosquitoes and represents as such a synapomorphy of this subgenus. *An. stephensi* males transfer steroid upon mating to females (Supplementary Fig. 2) as do *An. gambiae* males [13] and males of other *Cellia* species such as *Anopheles arabiensis* and *Anopheles dirus* [20]. This strongly suggests that transfer of steroids to females during mating is part of the “male 20E production” synapomorphy of *Cellia* mosquitoes. Our geographical mapping of nearly all contemporary mosquito species belonging to the *Cellia* subgenus (224 species) against the ones of *Anopheles* subgenus (184 species) (Figure 2) suggests that the common ancestor of the *Cellia* subgenus diverged after separation of South America and Africa in agreement with previous observations [3, 33]. This biogeographic calibration is consistent with divergence times obtained from our phylogenetic analysis placing the origin of steroid production by males of the *Cellia* subgenus around 84.1-69.2 Ma (112.7-45.6, 95% highest posterior density interval), *i.e*. after the separation of South America and Africa, which started at least 100 Ma ago with no land bridge for about 80 to 50 Ma [34, 35].

**Figure 2.**
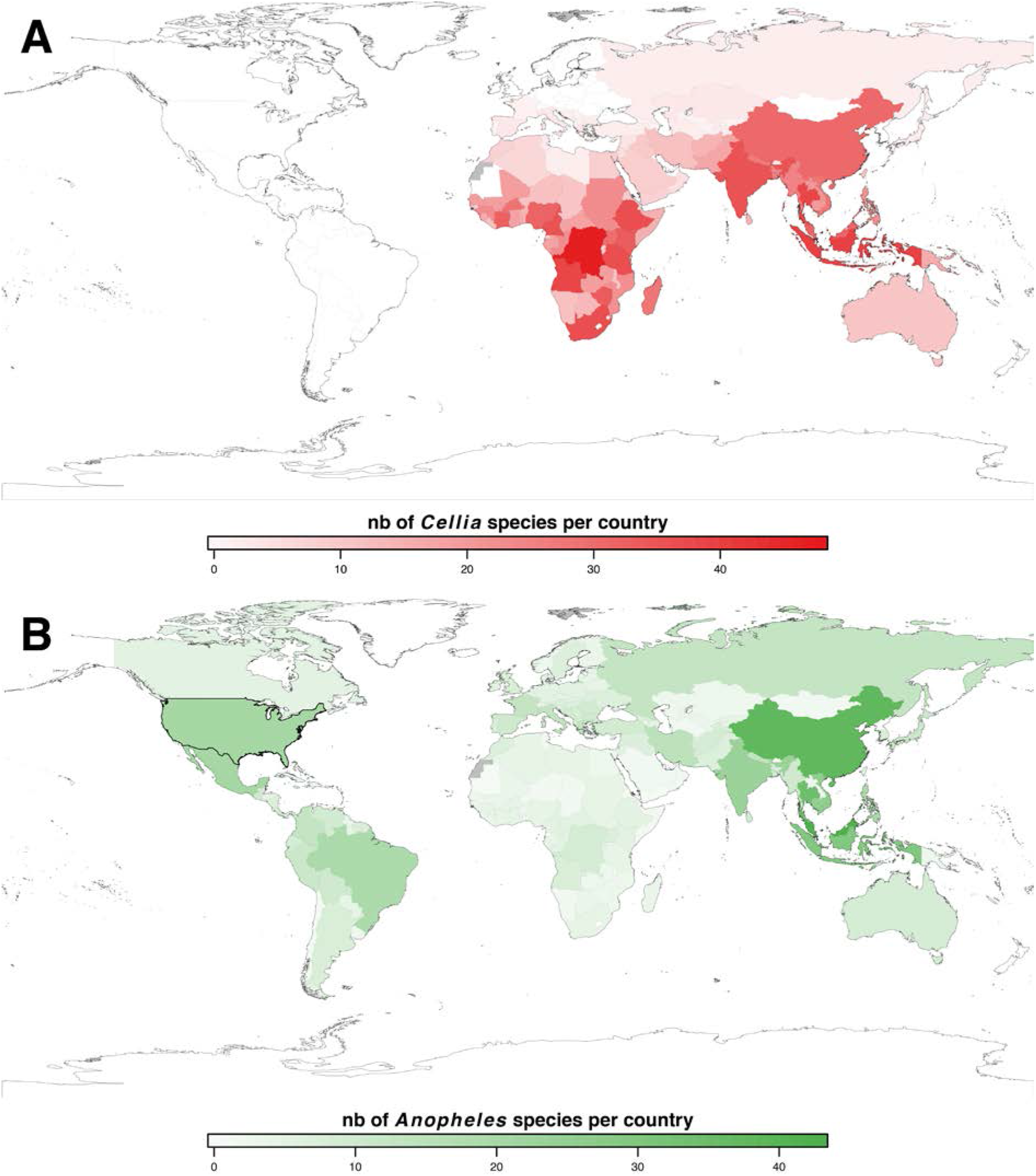
Present geographic distribution of species belonging to *Cellia* and *Anopheles* subgenera (*Anopheles* genus). Total numbers of mosquito species belonging to the *Cellia* (A, red) and *Anopheles* (B, green) subgenera per country (sourced from the Walter Reed Biosystematics Unit, http://www.wrbu.org/) are represented on world maps created with R. Numbers (nb) of mosquito species per country are represented by a coloured gradient as depicted under each map. Grey colour means no data are available for the country.

### Mating-induced effects do not correlate with sexually-transferred steroids

To get a deeper understanding of the effect(s) of sexually-transferred steroids in *Cellia* female mosquitoes, we investigated the influence of mating on two female reproductive traits which are known to be regulated by ovarian 20E. On the one hand, steroid hormones trigger ovarian follicle detachment from the germarium in *An. stephensi* and *Ae. aegypti* [14, 36], and ovarian steroids produced upon blood feeding stimulate vitellogenesis and egg development, on the other hand [14, 16–18, 37]. We therefore analysed these two phenotypes in virgin and mated females from 12 *Anopheles* species (8 *Cellia*, 3 *Anopheles* and 1 *Nyssorhynchus*). As expected, mating induces the separation of the secondary ovarian follicle from the germarium in *An. stephensi* (Figure 3, see also confocal pictures Supplementary Fig. 3), a *Cellia* species whose males produce and transfer steroids during mating. Mating induces as well the separation of the secondary follicle in *Anopheles minimus* and *Anopheles merus*, although to a lesser extent. However, no secondary follicle detachment was observed after mating in *An. dirus* nor detachment of the third follicle for the *Cellia* species whose secondary follicle is already fully detached in virgin females (*Anopheles farauti, An. gambiae, An. arabiensis* and *An. quadriannulatus*). Among the species whose males do not produce steroids, mating also induces secondary follicle detachment in some species (*Anopheles atroparvus* and *Anopheles freeborni*), but not in others, even in species showing partial detachment of the secondary follicle in virgin females *(Anopheles quadrimaculatus* and *Anopheles albimanus*). Similarly, mating increases the number of developed eggs in some mosquito species while not in others and this is not correlated to male steroid production (Figure 4 and Table 1). As an example, mating has an effect on egg development in *An. albimanus* (*Nyssorhynchus*), whose males do not produce steroids, but not in *An. gambiae* nor in *An. arabiensis* (*Cellia*). Intriguingly, those results are in discrepancy with previous studies reporting that mating increases egg development in *An. gambiae* and *An. arabiensis* [20, 38] but not in *An. albimanus* [20, 39]. These differences across studies are not due to the origin of the blood used to feed females (animal or human) as we obtained the same results with *An. albimanus* and *An. gambiae* fed on mouse or human blood (Supplementary Fig. 4). It is likely that some variations can be observed between strains of a single *Anopheles* species as already described for some *Aedes* species [40, 41]. It cannot be excluded that male steroids transferred upon mating benefit reproduction of some *Cellia* species but only under certain environmental conditions as different ecological pressures such as nutrition resources can favour or not the maintenance and importance of nuptial gifts, as shown for *Ae. aegypti* [42–44].

**Figure 3.**
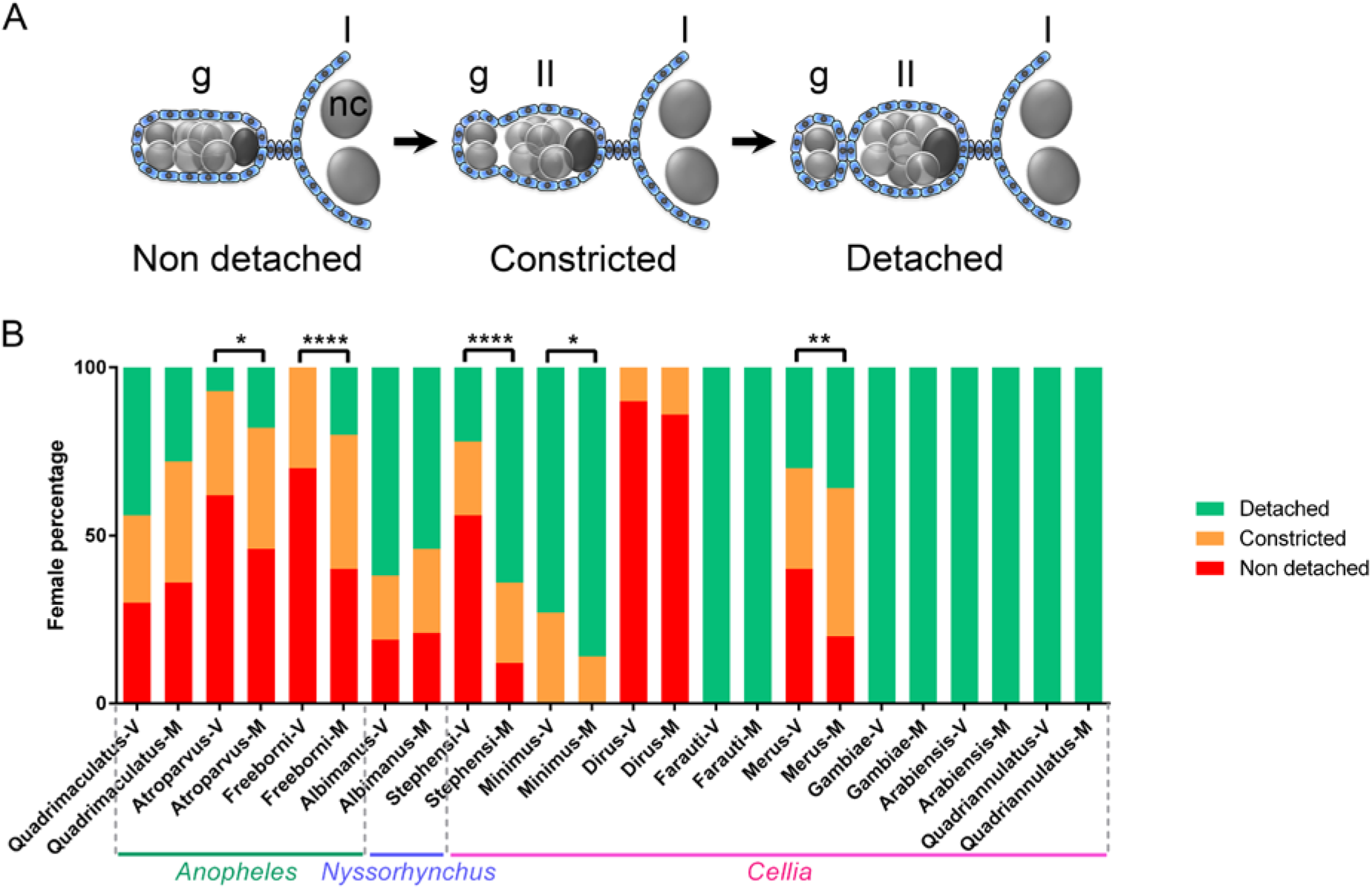
Secondary follicle detachment from the germarium in virgin and mated non blood-fed females from 12 *Anopheles species*. (A) Cartoon representing the detachment of the secondary follicle from the germarium. Follicular cells of somatic origin are coloured in blue. Germ cells (germline stem cells, developing cysts and nurse cells) are coloured in grey and the future oocyte of the secondary follicle in dark grey. g: germarium, I: primary follicle, II: secondary follicle, nc: nurse cells. (B) Secondary follicle detachment from the germarium in ovarioles of virgin (V) and mated (M) females. Secondary follicles are either detached from the germarium (detached, green), in the progress of detachment (constricted, yellow) or non-detached yet (non-detached, red). Representative pictures of these three states are shown in Supplementary Fig. 3. The secondary follicle are significantly more detached in mated females compared to virgin females for *An. atroparvus* (p=0.0226), *An. freeborni* (p<0.0001), *An. stephensi* (p<0.0001), *An. minimus* (p=0.0228) and *An. merus* (p=0.0072).

**Figure 4.**
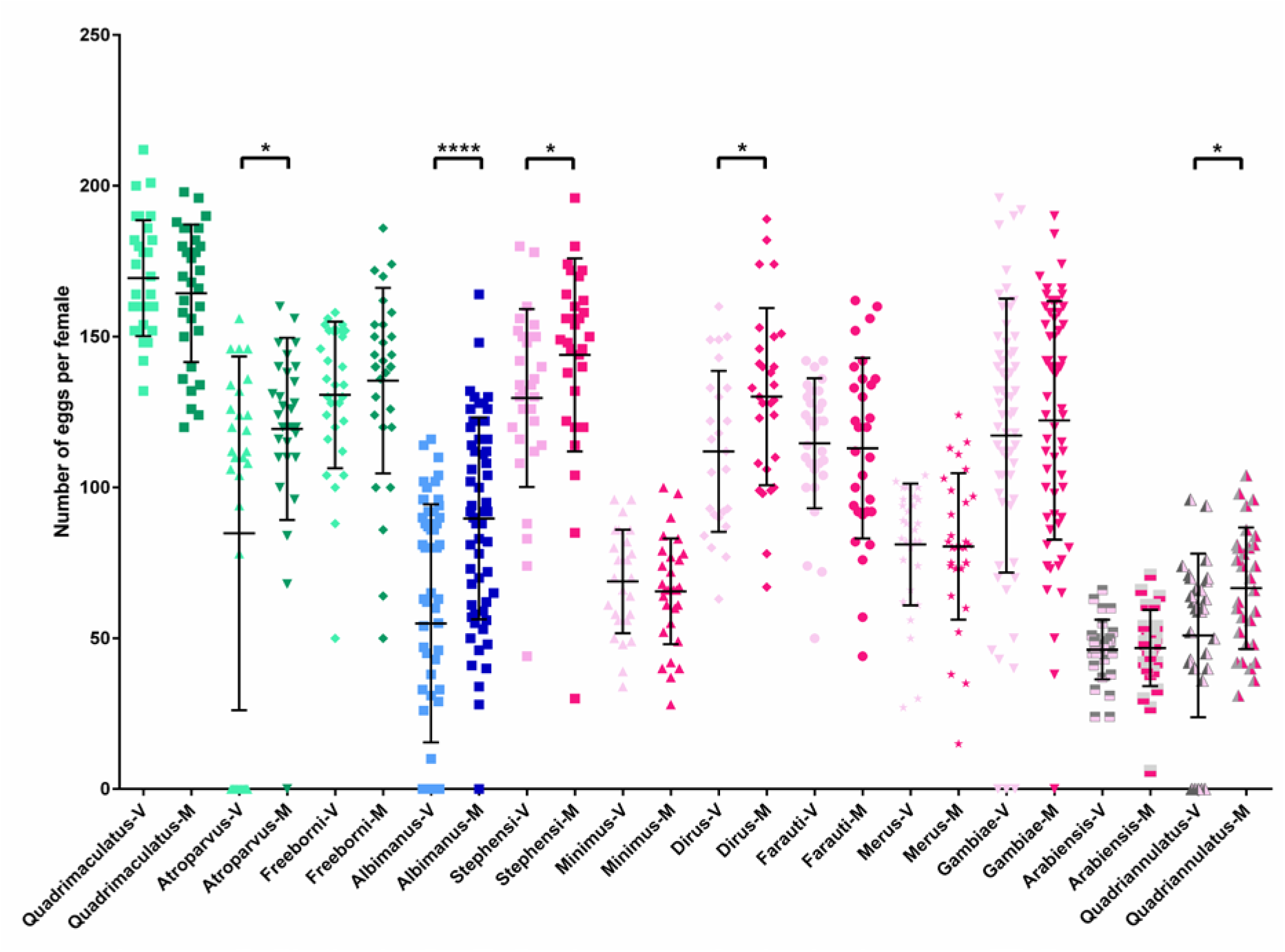
Egg development in virgin and mated blood-fed females from 12 species of *Anopheles* mosquitoes. Total number of eggs in virgin (V, light colours) and mated (M, dark colours) females 48 hours after blood feeding. Green: *Anopheles* subgenus, blue: *Nyssorhynchus* subgenus, pink: *Cellia* subgenus. Females from *An. atroparvus* (Mann-Whitney U=295.5, p= 0.0214), *An. albimanus* (Mann-Whitney U= 941.5, p<0.0001), *An. stephensi* (Mann-Whitney U= 232.5, p= 0.0235), *An. dirus* (Mann-Whitney U= 298, p= 0.0240) and *An. quadriannulatus* (Mann-Whitney U= 309.5, p= 0.0373) develop significantly more eggs when they are mated.

**Table 1.** Summary of mating effect on follicle detachment and egg development in regard to malaria vector status and male steroid production in *Anopheles* species. Dominant vector species (DVS) of human malaria are signalled by a +. For male steroid production, relative low titers (mean range 500pg E equivalent per male) are indicated by +, medium (range [1000-2000] pg E equivalent per male) titers by ++, and high titers (above 3000pg E equivalent per male) by +++. For follicle detachment and increase of egg development, - indicates no effect of mating and + indicates an effect of mating on either reproductive traits in females.

| Subgenus | Species | DVS | Male steroid<br>production | Follicle<br>detachment | Egg<br>development |
| --- | --- | --- | --- | --- | --- |
| *Anopheles* | *An. quadrimaculatus* | - | - | - | - |
|  | *An. atroparvus* | + | - | + | + |
|  | *An. freeborni* | + | - | + | - |
| *Nyssorhynchus* | *An. albimanus* | + | - | - | + |
| *Cellia* | *An. stephensi* | + | ++ | + | + |
|  | *An. minimus* | + | + | + | - |
|  | *An. dirus* | + | + | - | + |
|  | *An. farauti* | + | ++ | - | - |
|  | *An. merus* | + | + | + | - |
|  | *An. gambiae* | + | +++ | - | - |
|  | *An. arabiensis* | + | +++ | - | - |
|  | *An. quadriannulatus* | + | ++ | - | + |

Importantly, across the species investigated, mating induces detachment of the secondary follicle from the germarium in some species of which a low proportion of virgin females has the secondary follicle already detached as well as in some species of which a high proportion of virgin females has a detached follicle. Similarly, the occurrence of mating-induced effects on egg development is not linked to the ability of virgin females to develop a low or high number of eggs after blood feeding. Moreover, among species within the *Cellia* subgenus, the occurrence or absence of mating-induced phenotypes in females are not linked to the different quantities of steroids produced by the cognate males (Figure 1 and Table 1) and likely transferred to females during mating as determined for *An. gambiae, An. stephensi, An. arabiensis and An. dirus* (Supplementary Fig. 2, [13, 20]). Indeed, as depicted in Table 1, mating triggers both secondary follicle detachment and a rise in the number of developed eggs in *An. stephensi*, or only an increase in egg development in *An. dirus*, while this does not hold for *An. gambiae* and *An. arabiensis*, two species whose males produce and transfer higher amounts of steroids than *An. stephensi* and *An. dirus*. Overall, our analysis demonstrates that post-mating responses increasing fecundity in females indeed exist in *Cellia* species, some of which are likely mediated by male steroids. Importantly, they also occur in mosquito species whose males do not produce and transfer steroids such as in species belonging to the *Anopheles* and *Nyssorhynchus* subgenera (Table 1). Recently, male 20E has been shown to induce refractoriness to further mating in *An. gambiae* mosquitoes [19]. Similarly to results obtained for the two post-mating responses analysed in this study, female monoandry (insemination by a single male) occurs in species whose males produce steroids but also in ones whose males do not [45–48]. While few data are available for mosquitoes, it is well known that post-mating responses are quite conserved among insects. However, the rapid evolution of insect reproductive systems often results in species-specific genes and signalling pathways that ultimately trigger similar post-mating changes in different insect species [49]. For instance, *Ae. aegypti* males transfer to females Juvenile Hormone (JH) and *Drosophila* flies transfer male accessory gland peptides such as Sex Peptide upon copulation to trigger physiological and behavioural changes in mated females [50, 51]. Likewise, males from *Cellia* species transfer steroid hormones while males from *Anopheles* and *Nyssorhynchus* species are likely to transfer other and not yet identified molecule(s) to achieve similar effects. As JH is transferred to female during mating in *Ae. aegypti* mosquitoes but also in the Lepidoptera *Heliothis virescens* [52], sexual transfer of steroid hormones in other mosquito subgenera, genera or even insect orders not yet tested cannot even be excluded.

## Conclusions

A deep understanding of the selective forces driving reproductive strategy diversity and their functional consequences are critical for designing strategies for management of insect pests. It was previously suggested that post-mating responses in *Anopheles* mosquitoes only exist in species whose males produce and transfer steroids to females and that would be specific to dominant human malaria vectors [20]. Analysing the evolution of steroid production by male mosquitoes from a larger set of mosquito species reveals that this physiological trait is a synapomorphy of the *Cellia* subgenus. Importantly, there is no correlation between the evolution of sexually-transferred 20E and malaria transmission to humans (Table 1). Consistent with this, while we show that male steroid production and subsequent transfer to females is likely to have evolved only once in the common ancestor of *Cellia* species, phylogenetic analyses on malaria mosquitoes support a convergent evolution with independent acquisitions of vectorial capacities in *Anopheles* mosquitoes [53–55]. Furthermore, we demonstrate that mating-induced phenotypes are variable among species and possibly even among strains or under different environmental conditions. These differences are independent of males ability to produce and transfer steroids to females and are not correlated to malaria vectorial capacity. Apart from follicle detachment, increase in egg development and induction of refractoriness to mating, numerous other functions of steroids have been described in adult insects [19, 20, 56–61]. Thus, sexually-transferred steroids could mediate different functions with more or less direct benefit for the reproduction of *Cellia* females, due to the rapid evolution of reproductive systems between species. The recent interest in sexually-transferred steroids in anopheline mosquitoes has led to propose targeting 20E pathways with 20E receptor agonists to manipulate mosquito female reproduction as a mean to reduce vector populations and malaria transmission [21, 62]. In the light of our results, manipulating sexual-transfer of steroids in mosquitoes or using 20E receptor agonists in the field to manipulate *Anopheles* mosquito population and malaria transmission should be considered with caution in order to benefit malaria vector control strategies and also because it risks to affect other untargeted non-malaria vector species.

It remains open as what were the evolutionary forces that have initially promoted the acquisition and radiation of this presumably costly male steroid production and sexual gift to females in *Cellia* mosquitoes around 84.1-69.2 Ma. At this time, two main paleogeological events that had impact on environmental conditions may have led to pressures driving the evolution of steroid production and transfer by males in *Cellia* species: i) the Gondwana break up at 100 Ma with separation of South America and Africa [63]; ii) the Cretaceous-Paleogene extinction event at 65.5 Ma [64, 65]. Because such geographical isolation and environmental stresses are believed to drive traits contributing to animal species survival, it is likely that transfer of steroids by males to females have favoured *Cellia* species populations at this critical time.

## Methods

### Mosquito species and rearing

Ecdysteroid production by sexually mature males was analysed in 19 mosquito species. Sixteen (16) species belong to the subfamily *Anophelinae* (*An. arabiensis, An. dirus, An. farauti, An. gambiae* form M, *An. merus, An. minimus, An. quadriannulatus, An. stephensi, An. albimanus, An. albitarsis, An. aquasalis, An. atroparvus, An. freeborni, An. plumbeus, An. pseudopunctipennis* and *An. quadrimaculatus*) and 3 species belong to the subfamily *Culicinae (Ae. aegypti, Ae. albopictus* and *Cx pipiens*). *An. gambiae* form M, now called *An. coluzzii* (N’Gousso strain), *An. stephensi* (Sda 500) were permanently reared at Institut Pasteur (France). *An. albimanus* STECLA (MRA-126), *An. arabiensis* DONGOLA (males generously prepared by T. Bukhari and MRA-856), *An. quadrimaculatus* ORLANDO (MRA-139), *An. minimus* MINIMUS1 (MRA-729), *An. dirus* WRAIR2 (MRA-700), *An. farauti* FAR1 (MRA-489), *An. freeborni* F1 (MRA-130), *An. atroparvus* EBRO (MRA-493), *An. quadriannulatus* SANGWE (MRA-1155) and *An. merus* OPHANSI (MRA-803), were obtained through BEI Resources, NIAID, NIH, and contributed by M.Q Benedict (MRA-126,-856,-139,-729,-700,-489,-130,-493), C. Aranda (MRA-493), W. Takken (MRA-1155) and R. Mahraj (MRA-803). *Ae. aegypti* (Liverpool strain) and *Ae. albopictus* (Ho Chi Min Ville, Vietnam) were a generous gift from A-B. Failloux (Institut Pasteur, France). Eggs from *Cx. pipiens* (anautogenous strain) were kindly provided by M. Weill (ISEM, Montpellier, France). Mosquito larvae were reared at 27°C in deionized water supplemented with minerals and fed on TetraMin Baby-E fish food from the day of hatching to the fourth larval instar supplemented with pieces of cat food. Male and female adults were maintained at 27°C, under 68% relative humidity and a 12/12h light/dark cycle, and provided free access to a 10% wt/vol sucrose solution for the first 5 days post-emergence (PE). Female mosquitoes (first gonotrophic cycle) were allowed to feed for 30 min on the blood of an anesthetized mouse or rabbit depending on mosquito species preference (see Supplementary Table 2). *An. plumbeus* L4 larvae and pupae were harvested from a natural pond in Switzerland and reared up to the adult stage at the Institut Pasteur. Mature males of *An. albitarsis* and *An. aquasalis* were kindly prepared and provided by D. Valle and L. Moreira (Fiocruz, Brazil) and *An. pseudopunctipennis* by F. Lardeux (IRD, Bolivia).

### Mosquito sampling, ecdysteroid extraction and quantification

To measure ecysteroid titers in virgin adult males and females, males and females were separated on the day of adult emergence, transferred individually 5 days later in methanol and stored at −20°C until ecdysteroid extraction. For the transfer experiments, *An. stephensi* males and females were separated on the day of adult emergence for 5 days to allow male sexual maturation. Mating experiments and sampling were performed as described in [13]. Total ecdysteroids from individual whole mosquitoes were extracted with methanol and re-dissolved in enzyme immuno assay (EIA) buffer. Empty tubes were treated similarly in parallel to be used as a negative control (referred as extraction blank). Ecdysteroids were quantified by EIA, with 20-hydroxyecdysone-2-succinate coupled to peroxydase as a tracer (dilution 1:100,000) and the L2 antiserum (gift from M. De Reggi (Marseille, France); dilution 1:100,000). Calibration curves were generated with ecdysone (E; 3,6 - 500 pg/tube) diluted in EIA buffer, and titers were expressed as E equivalents. Under these conditions, detection limit is 2 pg E equivalents. All measurements were performed in duplicate and the results are expressed as mean values ±SEM of several (n=20 at least) independent samples and have been repeated on two independent cohorts of mosquitoes. Samples at or above the highest value of the calibration curve were diluted and quantify again. The intra- and inter-assay variation coefficients were 3,9% and 5,6%, respectively. For steroid titers in whole males from different species, data were subjected to statistical analysis using Kruskall-Wallis test for nonparametric data followed by Dunn’s post-test (control group: extraction blank). The indicated p values are those obtained with Dunn’s test. For the transfer experiment in *An. stephensi*, results were subjected to statistical analysis using Mann-Whitney test.

### Taxon sampling and DNA sequencing for phylogenetic analysis

We chose 20 *Culicidae* species for phylogenetic and comparative analysis. We selected 16 species of the genus *Anopheles* (*Anophelinae* subfamily) of which *An. arabiensis, An. dirus, An. farauti, An. gambiae, An. merus, An. minimus, An. stephensi, An. quadriannulatus, An. atroparvus, An. freeborni, An. plumbeus, An. pseudopunctipennis, An. quadrimaculatus, An. albimanus, An. albitarsis*, and *An. aquasalis*. As outgroups, we chose *Chagasia bathana* (*Anophelinae* subfamily, *Chagasia* genus), and 3 mosquito species belonging to the subfamily *Culicinae* with *Ae. aegypti, Ae. albopictus* (*Aedes* genus) and *Cx. pipiens (Culex* genus). Sequence data were generated for *An. aquasalis*, *An. atroparvus*, *An. merus* and *An. plumbeus*. Genomic DNA was obtained from single individuals using the DNeasy Blood and tissue kit (QIAGEN). A common set of molecular markers were chosen based on the availability of sequence data for *Ch. bathana*. Partial genomic regions of four nuclear genes (*g6pd, white, 18S and 28S)* and four mitochondrial genes (*COI, COII, ND5* and *CYTB*) were amplified by PCR with gene-specific or degenerate primers (sequences in Supplementary Table 3). For PCR amplifications, we used 0.4 μM oligonucleotides, 1 U GoTaq^®^ DNA Polymerase (Promega) per 35 μl reaction volume, 2 mM MgCl_2_, and 200 μM dNTP and reactions were carried out using standard thermocycle conditions. PCR products were purified and Sanger-sequenced with gene-specific primers or with T7, SP6 universal primers at Cogenics (www.cogenics.com, Beckman Coulter, GenBank Accession Numbers in Supplementary Table 4). Sequence data of the remaining species were obtained from GenBank and VectorBase (Supplementary Table 4). Sequences were examined and aligned with Geneious 6.1.3 (Biomatters). Our dataset was not complete, we did not find or generate sequence data for 10 / 160 (8 genes X 20 species) gene specific sequences and among the rest 15 / 150 of sequences were only partially covering the locus. The missing sequence data represent 5.6% of the total dataset and was annotated as a “?” (missing value) in the alignments for phylogenetic analysis. Introns were removed from *g6pd* and *white* sequences and the extremities of all protein coding sequences were trimmed to be in codon frame. Alignments for protein coding genes were re-aligned with the Geneious translation alignment program. In addition to gene specific alignments a concatenated dataset was generated in the following order: *COI-COII-ND5-CYTB-18S-28S-g6pd-white*. The number of informative sites was calculated using MEGA4 [66].

### Phylogenetic analysis

We used DNA sequences of partial regions of the coding sequence of mitochondrial genes (*COI, COII, ND5 and CYTB*) and nuclear ones (*18S, 28S, g6pd* and *white*) from the 19 mosquito species plus *Chagasia bathana*. The combined dataset had 4398 positions, including 2602 constant positions, 1776 variable positions and 1356 parsimony informative positions. Phylogenetic analysis of the concatenated, five-partition data set was performed by maximum likelihood in PhyML [67] and by Bayesian inference in BEAST [68]. For all analyses, partition specific models of nucleotide substitution were selected using the Akaike Information Criterion as calculated in jModelTest 2.1.3 [69, 70]. Maximum likelihood inference was done on the concatenated dataset in PhyML [67] using a GTR+I+G model of nucleotide substitution. Node support was determined by performing 100 bootstrap replicates. Bayesian phylogenetic analysis was performed with BEAST v1.7.5 [68] on the concatenated data set using five partitions with unlinked models of nucleotide substitution (Supplementary Table 5). The partitions corresponded to a single partition for mitochondrial sequences (*COI, COII, ND5* and *CYTB*) and four more partitions for the genes *18S, 28S, g6pd*, and *white*. Mitochondrial genes were combined into one partition because they are closely linked in the mitochondrial genome and largely evolve as a single unit with little to no recombination [71]. We used a common strict clock model and a yule birth process as tree prior. While the Bayesian approach places the species of the subgenus *Nyssorhynchus* as sister group of the *Anopheles* subgenus species, they formed the outgroup of the *Anopheles* and *Cellia* lineages in the maximum likelihood approach. Overall, the Bayesian phylogeny revealed high posterior probabilities for each node (>0.9) while the maximum likelihood analysis lacked strong statistical support at the nodes that separate the lineages of the three *Anopheles* subgenera. For species divergence time estimates, we therefore used the Bayesian phylogenetic analysis. We assigned calibration fossil ages to set priors for most recent common ancestors (MRCA). We used *Cx. winchesteri* (33,9-55,8 Ma) [72] (see also http://mosquito-taxonomic-inventory.info/category/fossil-culicidae/fossil-culicidae) to approximate the age of the most recent node shared by *Cx. pipiens* – and *Ae. aegypti/ Ae. albopictus*; and *An. dominicanus* (33,9-40,4 Ma) [73, 74] to estimate the age of the most recent node shared by *Anopheles* species. MRCA priors that incorporate fossil calibration dates were assumed to follow an exponential distribution, in the two above mentioned cases with an offset of 33.9 Ma and a mean according to the mean ages of the fossils. Based on a recently published phylogeny of Kamali *et al*. [53], the root age of the *Culicidae* was set to 147 Ma and the prior was assumed to be normally distributed with a standard deviation of 20 Ma. The Markov-Chain Monte-Carlo (MCMC) run was performed with a chain length of 10^8^ generations and was recorded every 1000 generations. Estimates were computed with Tracer version 1.5 (http://tree.bio.ed.ac.uk/software/tracer/) and MCMC output analysis was done using TreeAnnotator [68]. The first 2000 sampled trees were discarded as the burn-in. The phylogeny was visualized and annotated with Figtree version 1.4 (http://tree.bio.ed.ac.uk/software/figtree/).

### *Cellia* and *Anopheles* species distribution mapping

Total numbers of mosquito species belonging to the *Cellia* or *Anopheles* subgenera per country were taken from the Walter Reed Biosystematics Unit (WRBU, http://www.wrbu.org/). Data were further represented on maps created with R [75] using the “sp” package [76].

### Analysis of the secondary follicle detachment in virgin and mated non-blood fed females

Males and females were separated upon emergence to keep females virgin. A portion of females were put in a cage with males to obtain mated females. After 7 days, female ovaries were dissected, and the spermatheca of mated females checked for presence of spermatozoa. Ovaries from each female were then mounted on a slide in mounting media and observation of the secondary follicle detachment was performed using a transmitted light microscope. 30 to 60 females were analysed per species and per condition. Data were subjected to Chi-square test. For *An. stephensi*, ovaries from either virgin or mated females were fixed with 4% paraformaldehyde, washed 3 times with PBS-Tween 20 0.05% (PBS-T) and stained with DAPI. After 4 washes with PBS-T, ovaries were mounted on a slide in mounting media. Pictures were taken using a SP5 Leica confocal microscope.

### Analysis of egg development in virgin and mated blood fed females

Virgin and mated females were prepared as described above. A portion of females were put in a cage with males to obtain mated females. On 7 days PE, females were allowed to blood feed on anesthetized mouse or rabbit (Supplementary Table 2). *An. gambiae* and *An. albimanus* females were also fed with fresh human blood (ICAReB Platform, Center for Translational Research, Institut Pasteur, Paris, France) for comparison with mouse-fed mosquitoes. Ovaries were dissected 48 hours after blood feeding, and the total number of eggs in virgin and mated females was counted. Mated status was verified by observing spermatozoa in the spermatheca and only females with a filled spermatheca were taken into account. 30 to 60 females were dissected per species and per condition. Data were subjected to Mann-Whitney non-parametric test.

## Acknowledgments

We are grateful to L. Lambrechts, S. Blandin and J-P. Parvy for comments on the manuscript. We acknowledge D. Streicker, E. Hutchinson and M. Palmarini for discussions. We acknowledge M-T. Lecoq, S. Touron and C. Thouvenot for mosquito rearing, A-B. Failloux for providing *Ae. aegypti* and *Ae. albopictus* eggs, M. Weill (ISEM, Montpellier, France) for providing *Cx. pipiens* eggs, T. Bukhari for providing *An. arabiensis* samples, D. Valle and L. Moreira (FIOCRUZ, Brazil) for providing *An. aquasalis* and *An. albitarsis* samples, as well as F. Lardeux (IRD, Centre de Bolivie, La Paz, Bolivia) for providing *An. pseudopunctipenni*s samples. We thank E. Perret (Dynamic Imaging Platform, Institut Pasteur) for her assistance with confocal microscopy. The following reagents were obtained through BEI resources, NIAID, NIH: eggs of *An. albimanus* STECLA (MRA-126), *An. arabiensis* DONGOLA (MRA-856), *An. quadrimaculatus* ORLANDO (MRA-139), *An. quadriannulatus* SANGWE (MRA-1155), *An. merus* OPHANSI (MRA-803), *An. minimus* MINIMUS1 (MRA-729), *An. dirus* WRAIR2 (MRA-700), *An. farauti s.s*. FAR1 (MRA-489), *An. atroparvus* EBRO (MRA-493) and *An. freeborni* F1 (MRA-130).

## Data availability

The sequence data reported in this paper are tabulated in the Supplementary information and archived at Genbank.

## Contributions

E.P. and C.B. conceptualised and supervised the project as well as the experimental design; E.P., N.P., M.L., F.C., F.S., C.D-V. and E.B. performed the experiments; E.P., N.P., M.L., E.B. and C.B. contributed to data interpretation; E.P., N.P., M.L., F.S. and E.B. contribute to figure preparation; E.P. and C.B. wrote the manuscript; M.L., F.S., E.B. commented on the final manuscript.

## Ethical compliance

This study complied with all relevant ethical regulations. Project (n° 2013-0132) approved by the Ministère de l’Enseignement Supérieur et de la Recherche – Direction Générale pour la Recherche et l’Innovation – Secrétariat « Autorisation de projet » - 1, rue Descartes, 75231 PARIS cedex 5.

## Supplementary Figures and Tables

**Supplementary Figure 1.**
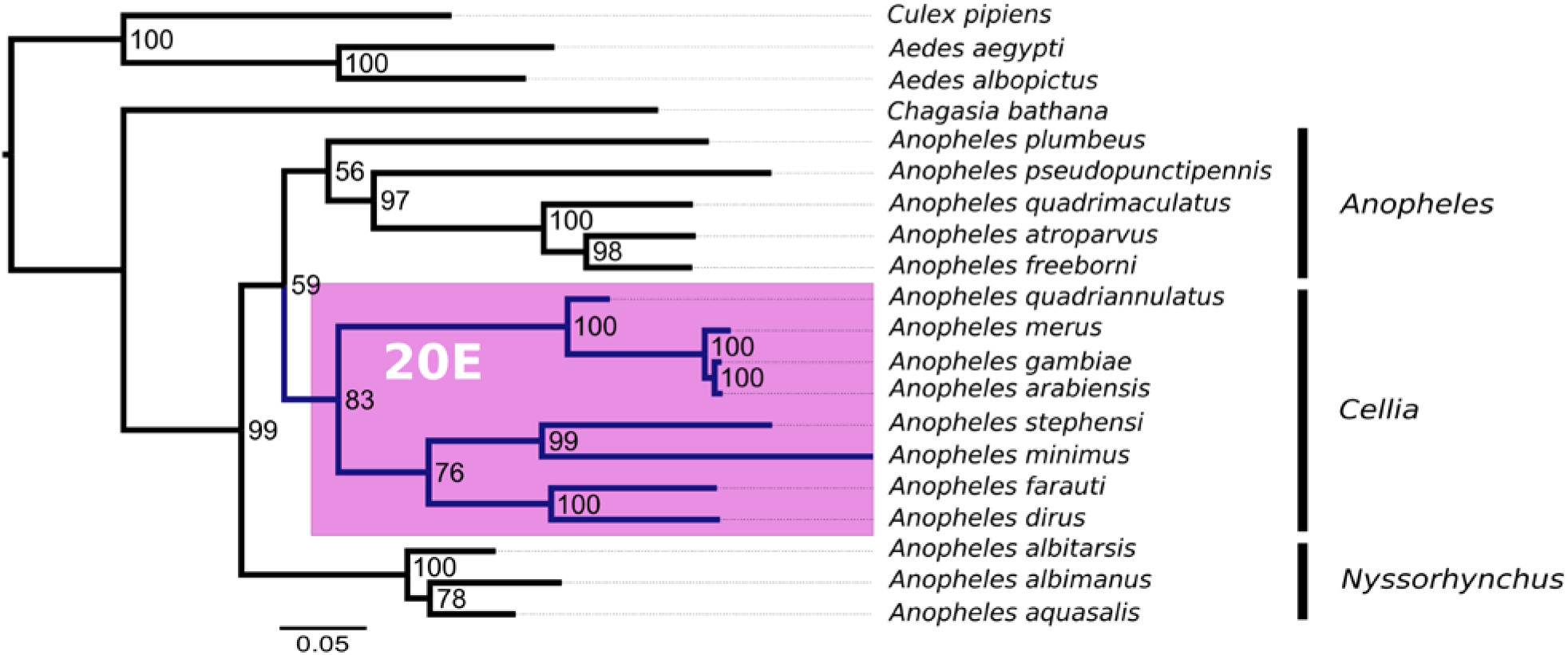
Phylogenetic relationships of the genus *Anopheles* and evolution of male 20E production. Maximum likelihood phylogeny (PhyML) based on a concatenated dataset. Bootstrap supports (100 replicates) are presented on the right side of each node. Bars on the right side indicate species that belong to the same subgenus. The lineages of the subgenus *Cellia* are highlighted in blue. The lineages with male 20E production are shaded in pink.

**Supplementary Figure 2.**
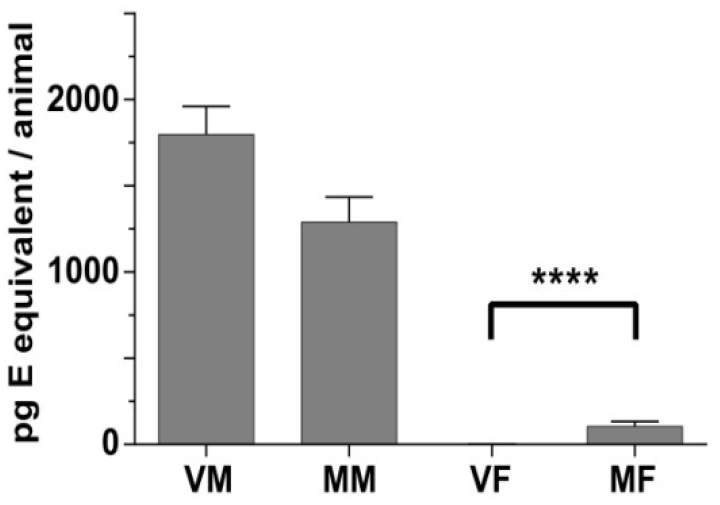
*Anopheles stephensi* males transfer ecdysteroids to females during mating. Ecdysteroid titers were measured in virgin males (VM), in mated males (MM) just after copulation, in virgin females (VF), and in mated females (MF) just after copulation. Ecdysteroids were extracted from each individual mosquito and quantified by EIA. Results are expressed as mean +/− SEM in in pg E equivalents per animal. Results were subjected to statistical analysis using Mann-Whitney test (****, P< 0.0001).

**Supplementary Figure 3.**
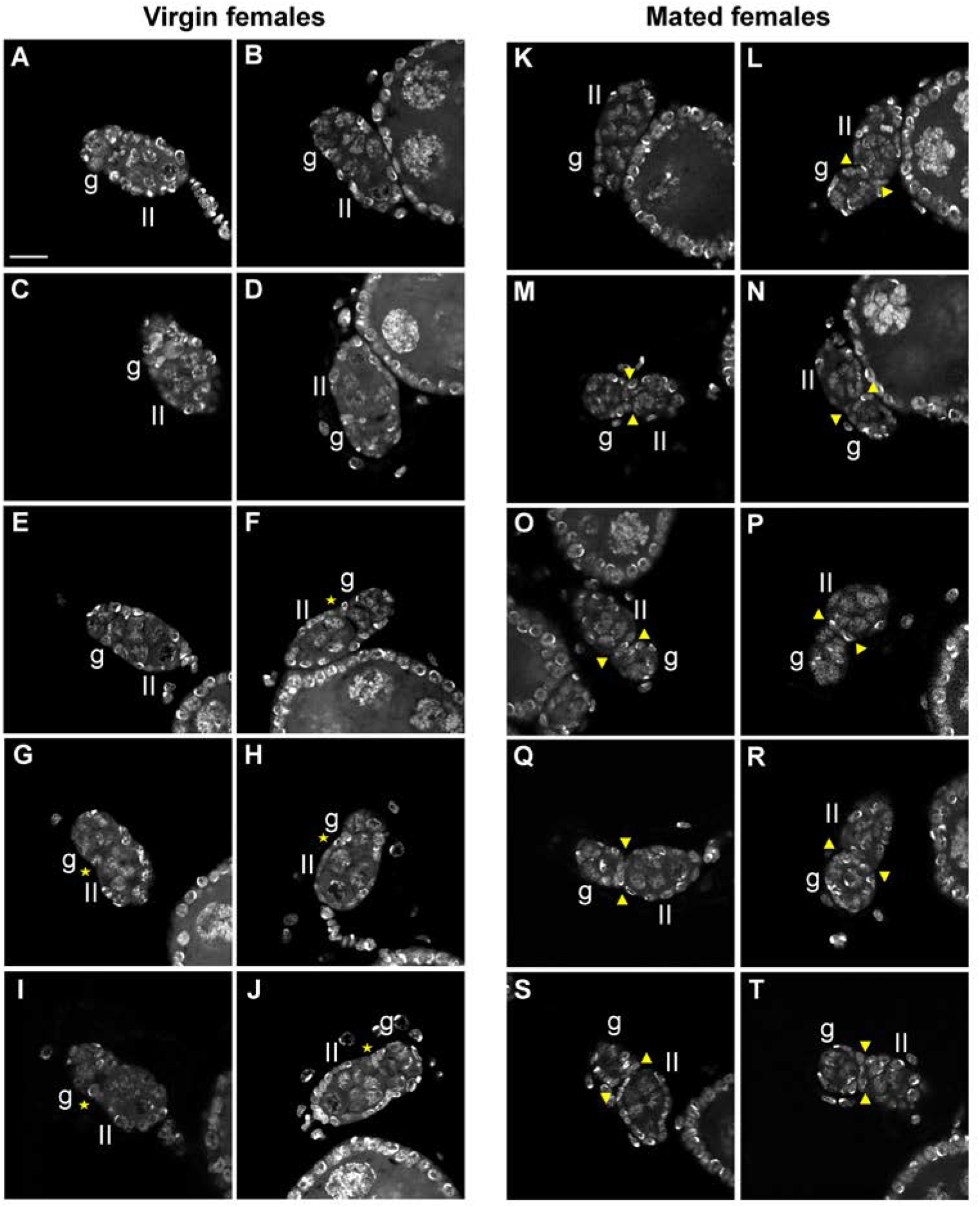
Secondary follicle detachment from the germarium in *Anopheles stephensi* non blood-fed females according to the insemination status. Confocal pictures of ovarioles showing the germarium and the secondary follicle from 10 non blood-fed females either virgin (A to J) or mated (K to T) stained with DAPI. g: germarium, II: secondary follicle. Yellow arrowheads show secondary follicles detached from the germarium and yellow stars show secondary follicles in progress of detachment. Follicles not marked are not detached. Scale bar is 10.25 μm.

**Supplementary Figure 4.**
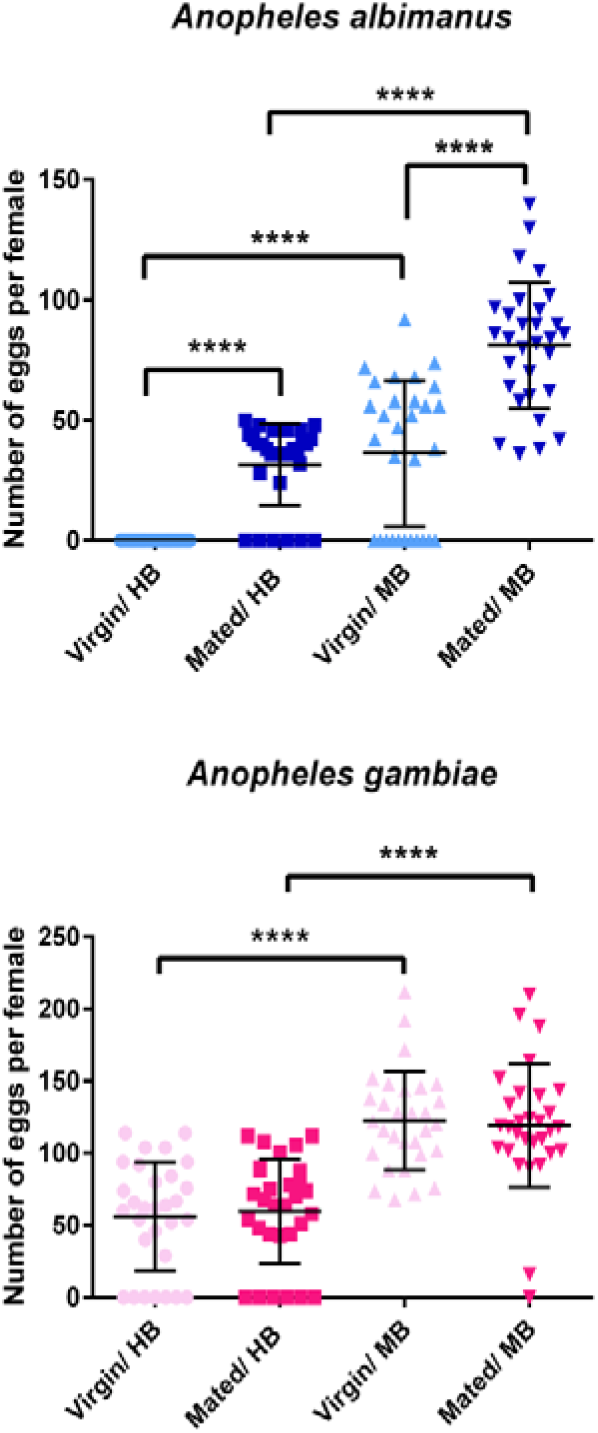
Egg development in *Anopheles albimanus* (*Nyssorhynchus* subgenus) and *Anopheles gambiae* (*Cellia* subgenus) virgin and mated females fed on mouse or human blood. Total number of eggs in virgin (light colours) and mated (dark colours) females was counted 48 hours after feeding on either human blood (HB) or mouse blood (MB). *An. albimanus* is coloured in blue and *An. gambiae* in pink. Data were subjected to Mann-Whitney non-parametric test. Mated females from *An. albimanus* develop significantly more eggs then virgin females when fed on human blood (Mann-Whitney U=90, p<0.0001) or mouse blood (Mann-Whitney U=122, p<0.0001). Virgin and mated females of both species develop significantly more eggs when fed on mouse blood compared to human blood (An. *albimanus* virgin: Mann-Whitney U=165, p<0.0001; *An. albimanus* mated: Mann-Whitney U=52.50, p<0.0001; *An. gambiae* virgin: Mann-Whitney U=74, p<0.0001; *An. gambiae* mated: Mann-Whitney U=90, p<0.0001).

**Supplementary Table 1.** Divergence time estimates of the *Anopheles* species and outgroups. Estimates were obtained from the Bayesian phylogenetic analysis and are based on fossil data for temporal calibration.

| Node | Taxon Set | Divergence Time* | Node | Taxon Set | Divergence Time* |
| --- | --- | --- | --- | --- | --- |
| 1 | all | 124.2 Ma (83.7-165.8) | 11 | *An. atroparvus* – *An. quadrimaculatus* | 25.6 Ma (16.7-34.8) |
| 2 | *Aedes* – *Culex*<br>(genus) | 78.9 Ma (51.6-106.5) | 12 | *An. atroparvus* – *An. freeborni* | 18.1 Ma (11.5-24.7) |
| 3 | *Aedes*<br>(genus) | 35.6 Ma (23.1-48.3) | 13 | *Cellia*<br>(subgenus) | 69.2 Ma (45.6-93.0) |
| 4 | *Anopheles* – *Chagasia*<br>(genus) | 113.7 Ma (75.2-152.4) | 14 | *An. farauti* – *An. stephensi* | 62.5 Ma (41.1-84.3) |
| 5 | *Anopheles*<br>(genus) | 84.1 Ma (55.8-112.7) | 15 | *An. stephensi* – *An. minimus* | 46.6 Ma (30.0-62.8) |
| 6 | *Nyssorhynchus* – *Anopheles*<br>(subgenus) | 76.4 Ma (50.4-102.3) | 16 | *An. farauti* – *An. dirus* | 31.6 Ma (20.5-42.9) |
| 7 | *Nyssorhynchus*<br>(subgenus) | 20.7 Ma (13.5-28.3) | 17 | *An. gambiae* – *An. quadriannulatus* | 17.2 Ma (11.0-23.7) |
| 8 | *An. aquasalis* – *An. albitarsis* | 17.4 Ma (11.1-24.1) | 18 | *An. gambiae* – *An. merus* | 3.2 Ma (1.9-4.5) |
| 9 | *Anopheles*<br>(subgenus) | 71.9 Ma (47.7-97.0) | 19 | *An. gambiae* – *An. arabiensis* | 1.2 Ma (0.6-1.9) |
| 10 | *An. atroparvus* – *An. pseudopunctipennis* | 61.9 Ma (40.3-82.9) |  |  |  |

\*Estimates are the posterior means with 95% highest posterior density intervals.

**Supplementary Table 2.** Animal species or blood source on which mosquitoes were fed in this study.

| ***Anopheles* species** | **Animal species** | **Blood source** |
| --- | --- | --- |
| *An. quadrimaculatus* | mouse |  |
| *An. atroparvus* | rabbit |  |
| *An. freeborni* | mouse |  |
| *An. albimanus* | mouse | human |
| *An. stephensi* | mouse |  |
| *An. minimus* | mouse |  |
| *An. dirus* | rabbit |  |
| *An. farauti* | mouse |  |
| *An. merus* | mouse |  |
| *An. gambiae* | mouse | human |
| *An. arabiensis* | rabbit |  |
| *An. quadriannulatus* | mouse |  |

**Supplementary Table 3.**
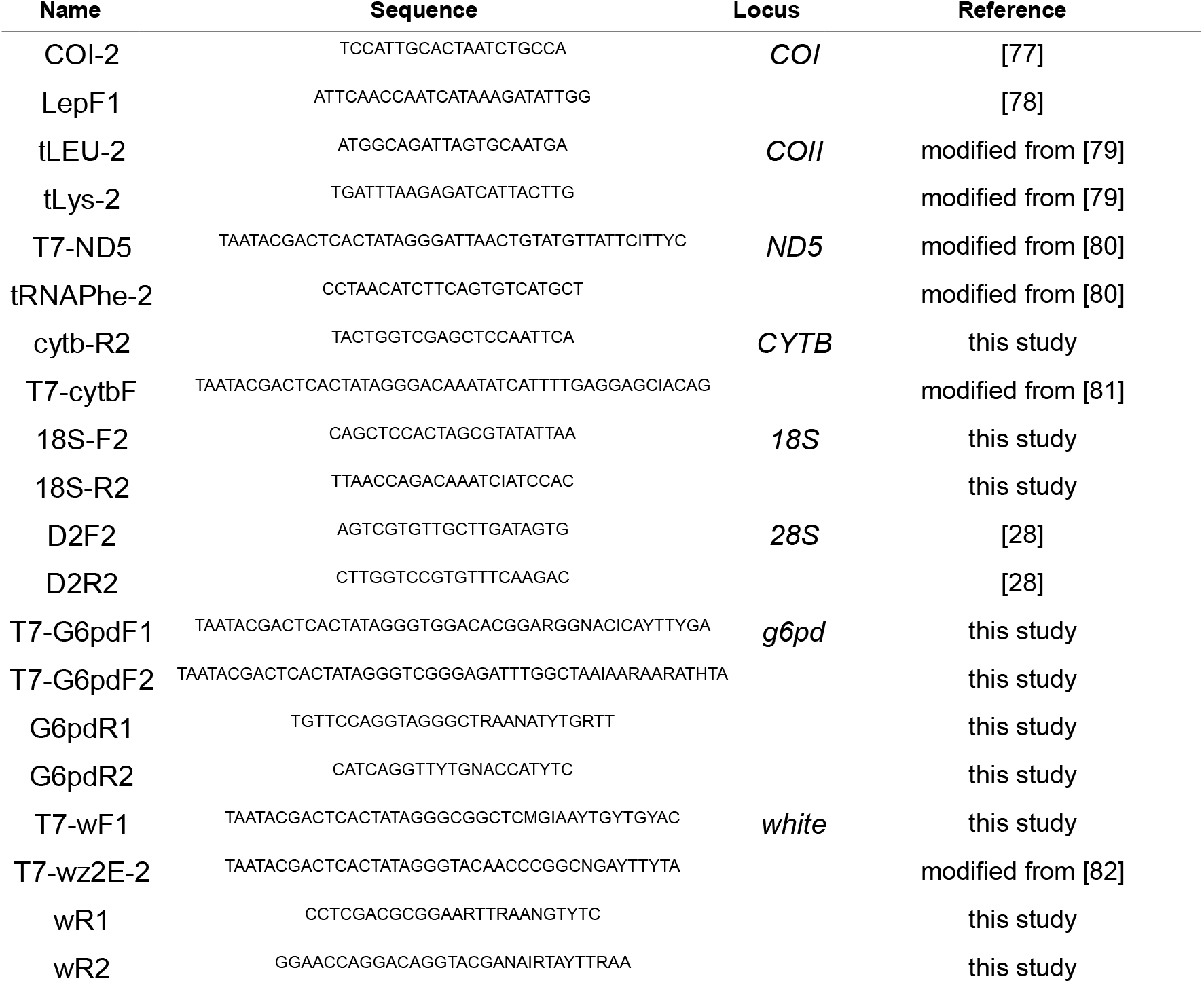
Sequences of primers used to sequence DNA for the phylogenetic analyses. Partial genomic regions of four nuclear genes (*g6pd*, *white, 18S* and *28S*) and four mitochondrial genes (*COI, COII, ND5* and *CYTB*) were amplified by PCR with gene-specific or degenerate primers. Degenerate oligonucleotides were designed based on previous studies and optionally modified or newly designed. Some oligonucleotide sequences contained T7 or SP6 universal primer sequences at their 5’ end, following Bonacum *et al*. [83].

**Supplementary Table 4.**
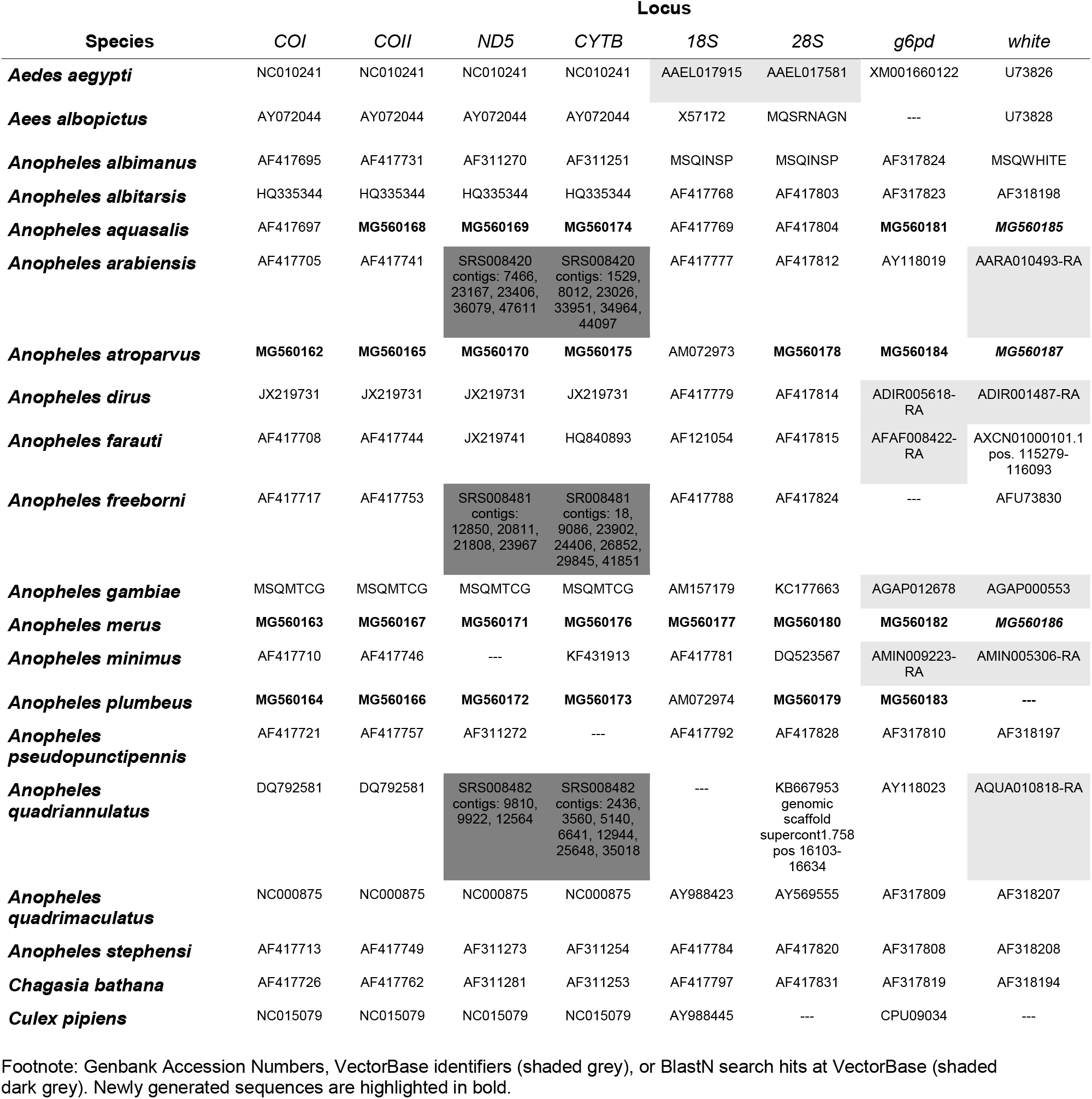
GenBank Accession Numbers of the dataset used for the phylogenetic analyses.

**Supplementary Table 5.**
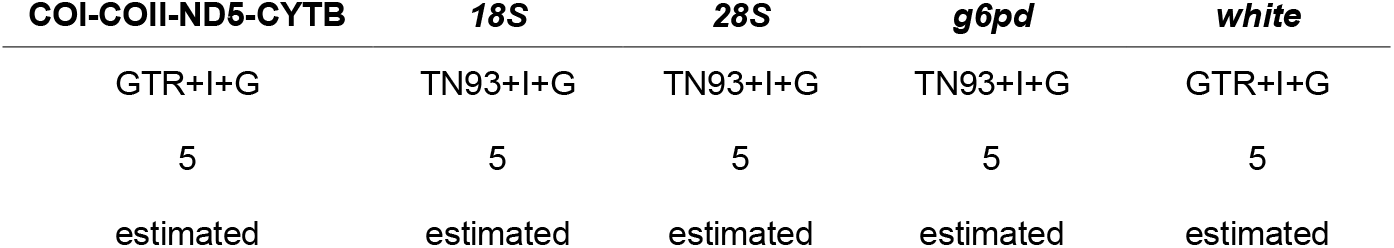
Partition specific site models and parameters.

